# MaXLinker: proteome-wide cross-link identifications with high specificity and sensitivity

**DOI:** 10.1101/526897

**Authors:** Kumar Yugandhar, Ting-Yi Wang, Alden King-Yung Leung, Michael Charles Lanz, Ievgen Motorykin, Jin Liang, Elnur Elyar Shayhidin, Marcus Bustamante Smolka, Sheng Zhang, Haiyuan Yu

## Abstract

Protein-protein interactions play a vital role in nearly all cellular functions. Hence, understanding their interaction patterns and three-dimensional structural conformations can provide crucial insights about various biological processes and underlying molecular mechanisms for many disease phenotypes. Cross-linking mass spectrometry has the unique capability to detect protein-protein interactions at a large scale along with spatial constraints between interaction partners. However, the current cross-link search algorithms follow an “MS2-centric” approach and, as a result, suffer from a high rate of mis-identified cross-links (~15%). We address this urgent problem, by designing a novel “MS3-centric” approach for cross-link identification and implemented it as a search engine called MaXLinker. MaXLinker significantly outperforms the current state of the art search engine with up to 18-fold lower false positive rate. Additionally, MaXLinker results in up to 31% more cross-links, demonstrating its superior sensitivity and specificity. Moreover, we performed proteome-wide cross-linking mass spectrometry using K562 cells. Employing MaXLinker, we unveiled the most comprehensive set of 9,319 unique cross-links at 1% false discovery rate, comprising 8,051 intraprotein and 1,268 interprotein cross-links. Finally, we experimentally validated the quality of a large number of novel interactions identified in our study, providing a conclusive evidence for MaXLinker’s robust performance.

## INTRODUCTION

In the post-genomic era, one of the main goals of systems biology is to determine the functions of all the proteins of various organisms. In the cell, most proteins function through interacting with other proteins. Therefore, generating interactome network models with high quality and coverage is a necessary step in the process of developing predictive models for protein functions at the scale of the whole cell^1^. Furthermore, structural information for protein-protein interactions can serve as a crucial prerequisite for understanding the mechanism of protein function^2^.

Rapid advancements in the respective fields of cross-linking and mass spectrometry lead to the development of a powerful technique known as cross-linking mass spectrometry (XL-MS)^3, 4^. XL-MS has been demonstrated to be an efficient technology to capture distance constraints, thereby providing crucial information to decipher the interaction partners and dynamics of protein-protein interactions^5^. Development of efficient MS-cleavable chemical cross-linkers such as disuccinimidyl sulfoxide (DSSO)^6^ expanded the applications of XL-MS ranging from studying individual functional complexes^7^ to discovering proteome-wide interactions by drastically minimizing the database search space. Recently, Liu *et al*^8^ demonstrated the high-throughput capability of XL-MS approach with the first-ever proteome-wide XL-MS study on HeLa cell lysate. They utilized CID-ETD toggle approach and identified a set of 1822 cross-links at 1%FDR employing XlinkX, a state-of-the-art search engine for XL-MS. The traditional ‘target-decoy’ approach for estimating false discovery rate (FDR) in peptide spectrum matches (PSMs) was adapted to estimate quality of the identified cross-links (each individual cross-link identification is also known as a Cross-link Spectrum Match (CSM)). Since then, rapid advancements in terms of technical capability have been reported by utilizing and combining multiple levels and types of fragmentation methods.

Most recently, Liu *et al*^9^ compared multiple fragmentation schemes, including CID-MS2, CID-MS2-ETD-MS2, CID-MS2-MS3 and CID-MS2-MS3-ETD-MS2 (a combination of CID-MS2-ETD-MS2 and CID-MS2-MS3 approaches) utilizing an updated version of XlinkX (XlinkX v2.0). Apart from CID-MS2, all other approaches combine spectra from multiple MS levels (MS2 and MS3) or from different types of energy fragmentations (CID and ETD) or both. Their analysis revealed that, the ensemble approach (i.e., CID-MS2-MS3-ETD-MS2) resulted in the highest number of cross-links, followed by CID-MS2-MS3, CID-MS2-ETD-MS2 and CID-MS2. Moreover, utilizing sequence information only from MS3 spectra (a subset of CID-MS2-MS3) for cross-link identification (‘MS3-Only’) resulted in the least number of crosslinks among all the approaches. Hence the study concluded CID-MS2-MS3-ETD-MS2 and MS3-Only approaches to be the most and least informative approaches, respectively. However, the study did not assess quality of different approaches at the given FDR cut-off using a rigorous comparative analysis.

In this study, we perform systematic and rigorous quality assessment across different XL-MS acquisition strategies, inspired by approaches widely-used in machine learning^1, 10^. Based on these analyses, we noted that XlinkX results in high number of mis-identifications. Therefore, we developed and validated a novel search algorithm named MaXLinker, which is based on an innovative “MS3-centric” approach, designed to efficiently eliminate incorrect cross-link candidates. At a 1% FDR, MaXLinker has an 18-fold lower rate of mis-identifications than XlinkX. With MaXLinker in hand, we performed a large-scale proteome-wide XL-MS study on K562 cell lysate, yielding the largest XL-MS data set to date. We further validated the cross-links using available three-dimensional structures and through a systematic experimental validation of novel interactions identified in our study.

## RESULTS

### Current MS2-centric cross-link search algorithms are limited in their sensitivity and specificity

When compared to traditional PSM searches, the identification of CSMs from a proteome-wide study is markedly more complex. This fact motivated us to thoroughly examine the MS2-centric algorithms^9, 11^ for processing proteome-wide XL-MS data sets. XlinkX is currently the most widely-used MS2-centric software (it is available as a node within Proteome Discoverer^9^). Thus, we performed a systematic quality comparison of cross-links generated by XlinkX using data from multiple XL-MS acquisition strategies described in Liu *et al*^9^. First, we obtained corresponding raw files for the three fragmentation schemes CID-MS2-ETD-MS2, CID-MS2-MS3 and CID-MS2-MS3-ETD-MS2 (through email request to Dr. Fan Liu). Then we performed cross-link search using XlinkX software (implemented in Proteome Discoverer 2.2) at 1% FDR with a concatenated database containing sequences from *E. coli* proteome (true search space) and *S. cerevisiae* (false search space). It is important to note that XlinkX by default, generates a *reversed* version of the input database and uses it as a decoy database to estimate FDR. As a next step, we compared the three fragmentation approaches in terms of the number of incorrect unique CSMs (CSMs with at least one peptide from the *S. cerevisiae* search space, i.e., mis-identifications). The aim of this search is to re-assess the quality of cross-links at 1% FDR, with expected fraction of incorrect CSMs involving unambiguous peptides from *S. cerevisiae* to be less than 1%. Surprisingly, the fraction of incorrect CSMs range from 14.8% to as high as 26.9% across the three acquisition strategies (**Fig. 1a**). Upon closer examination, we observed that among the three approaches, CID-MS2-MS3 showed significantly lower proportion of incorrect CSMs (14.8%) followed by CID-MS2-ETD-MS2 (25.1%), and CID-MS2-MS3-ETD-MS2 (26.9%) approaches. This analysis clearly indicates that the methodology implemented in XlinkX does not adequately evaluate the quality of the identified CSMs. Therefore, utilizing only the number of identifications for comparative evaluations^9^ might not yield accurate conclusions about the capability of different acquisition strategies. As the FDR filtering is typically performed at the redundant CSM level by the conventional cross-link search algorithms (i.e., before the processing step that results in a unique list of CSMs), we repeated the analysis at redundant CSM level and observed results consistent with what was found at the unique CSM level (**Supplementary Figure 1**).

**Figure 1.**
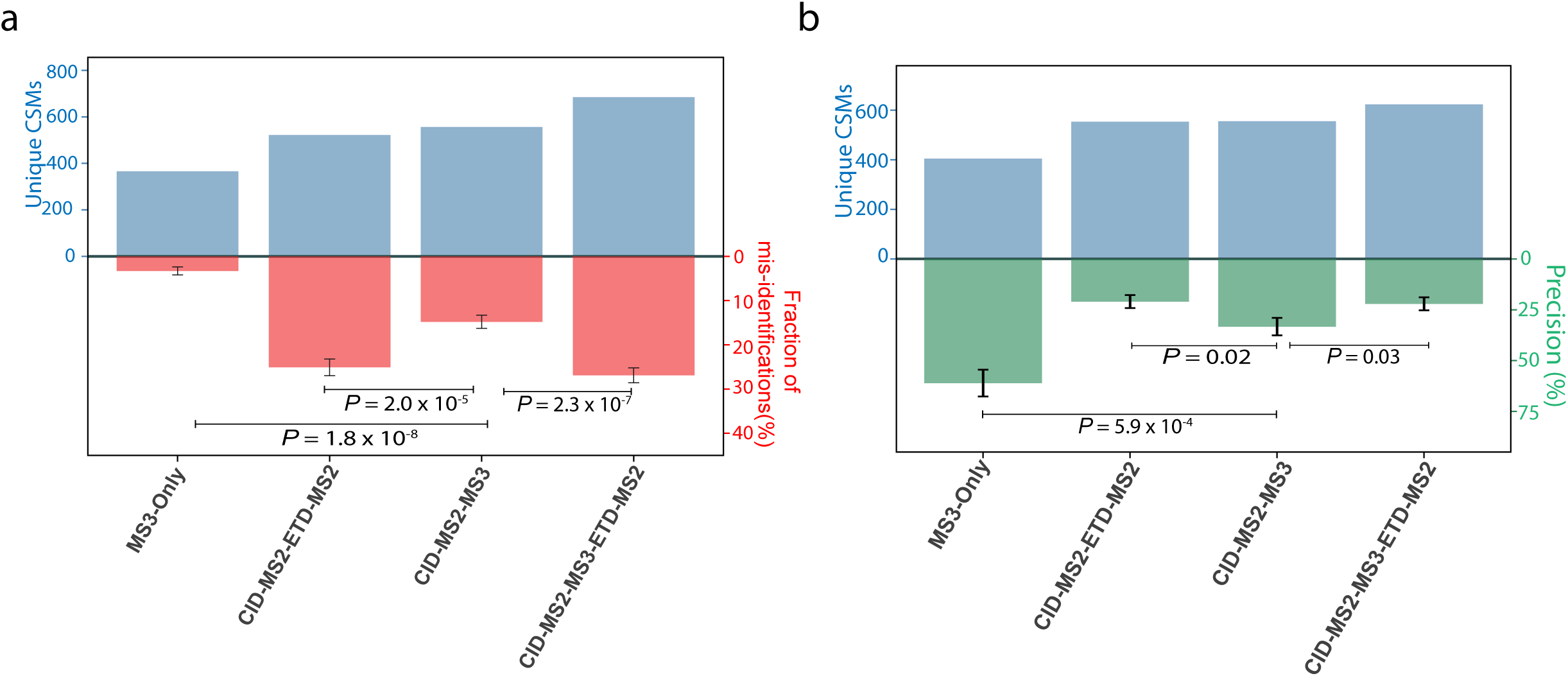
Comparative quality assessment between various acquisition methods for Cross-linking Mass-Spectrometry on six *E. coli* fractions from Liu *et al*^9^ (a) Comparison between different acquisition methods based on mis-identifications. (The search was performed using a database consisting amino acid sequences of *E. coli* and *S. cerevisiae* proteomes. Any CSM with either of the peptides exclusively from *S. cerevisiae* proteome was considered as a mis-identification). (b) Quality comparison across multiple acquisition methods using precision. A separate search was performed for panel ‘b’ using only the *E. coli* database in order to avoid underestimation of precision. (Significance was determined by a two-sided *Z*-test; The error bars represent the estimated standard error of mean)

### The most reliable sequence information for cross-linked peptides comes from the MS3-level

We also evaluated quality of identifications from CID-MS2-MS3 approach, with sequence information obtained exclusively from MS3 spectra (‘MS3-only’) (**Fig.1a**). Strikingly, we observed a drastically lower fraction of incorrect CSMs for ‘MS3-Only’ (3.3%), which is a subset of CID-MS2-MS3 (with 14.8% mis-identifications). This result clearly demonstrates that MS3, the most advanced MS level, provides higher quality sequence information in comparison to MS2-level. To improve the quality of XlinkX-identified CSMs, XlinkX allows the use of ‘Δ XlinkX score’ to further filter the CSMs. As a next step, we filtered the CSMs using five different ‘Δ XlinkX score’ cutoffs and re-assessed their quality across different approaches. We observed that, overall, increasing the stringency based on ‘Δ XlinkX score’ significantly reduced the number of incorrect CSMs for all three acquisition approaches (**Supplementary Figure 2**). However, even after filtering by ‘Δ XlinkX score’, the trend across the different methods was similar to what was observed before the filtering **(Fig.1a and Supplementary Figure 2**), with data from the MS3-level yielding the highest fraction of reliable CSMs.

### Precision is a reliable metric for comparative quality assessment for proteome-wide XL-MS data sets

To perform a more comprehensive and rigorous quality evaluation, we next utilized precision to compare the quality across the three acquisition approaches. Precision has been shown to be an effective quality measure in machine learning based studies for the identification of proteins^12, 13^ and their interactors^14^. Precision has also been utilized for evaluating the quality of large-scale interactions screens, where it is derived using known interactions (as *training set*)^1^. However, none of the reported XL-MS studies have adapted it as a quality estimate for their data sets. Here we present precision as a measure to assess quality of cross-link data sets from proteome-wide XL-MS studies. Precision for XL-MS essentially represents the fraction of identified interprotein cross-links that correspond to known protein-protein interactions (**METHODS**). Remarkably, precision complemented the result obtained in the above analysis using additional *S. cerevisiae* search space (**Fig. 1b, Supplementary Figure 2**).

**Figure 2.**
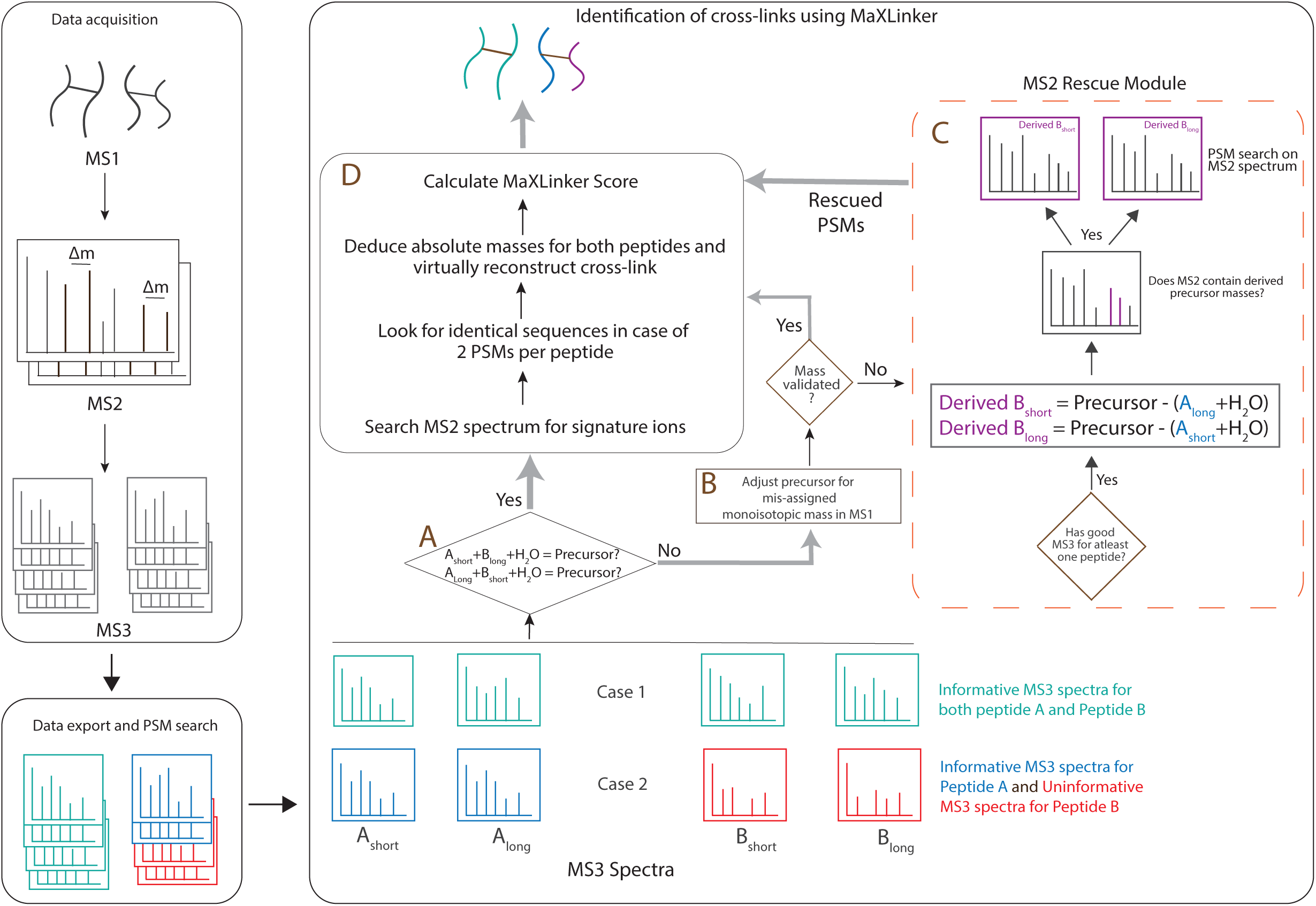
Overview of MaXLinker’s strategy for identification of cross-links from XL-MS

### Most of the reliable cross-link identifications are contributed by CID-MS2-MS3 methodology

It is important note that CID-MS2-MS3-ETD-MS2 (combination CID-MS2-MS3 and CID-MS2-ETD-MS2 methodologies) resulted in higher fraction of mis-identifications when compared to CID-MS2-MS3 approach (**Fig. 1a**). Upon closer examination of the quality of CSMs identified by the inherent CID-MS2-MS3 and CID-MS2-ETD-MS2 methodologies, we observed that at 1% FDR, CSMs identified exclusively by CID-MS2-ETD-MS2 contains almost two-fold higher fraction of mis-identifications in comparison to exclusive identifications by CID-MS2-MS3 (**Supplementary Figure 3**). We repeated the analysis after filtering the CSMs at different ‘Δ XlinkX score’ cut-offs. It is interesting to note that, as the cut-off score increases, the number of identifications contributed exclusively by CID-MS2-ETD-MS2 reduces consistently, to as low as 6% when compared to the exclusive identifications by CID-MS2-MS3 (at ‘Δ XlinkX score’ ≥ 50) (**Supplementary Figure 3**). These results reveal that, for CID-MS2-MS3-ETD-MS2, at higher quality cut-offs, CID-MS2-ETD-MS2 fails to yield additional cross-links than what were already captured by CID-MS2-MS3.

**Figure 3.**
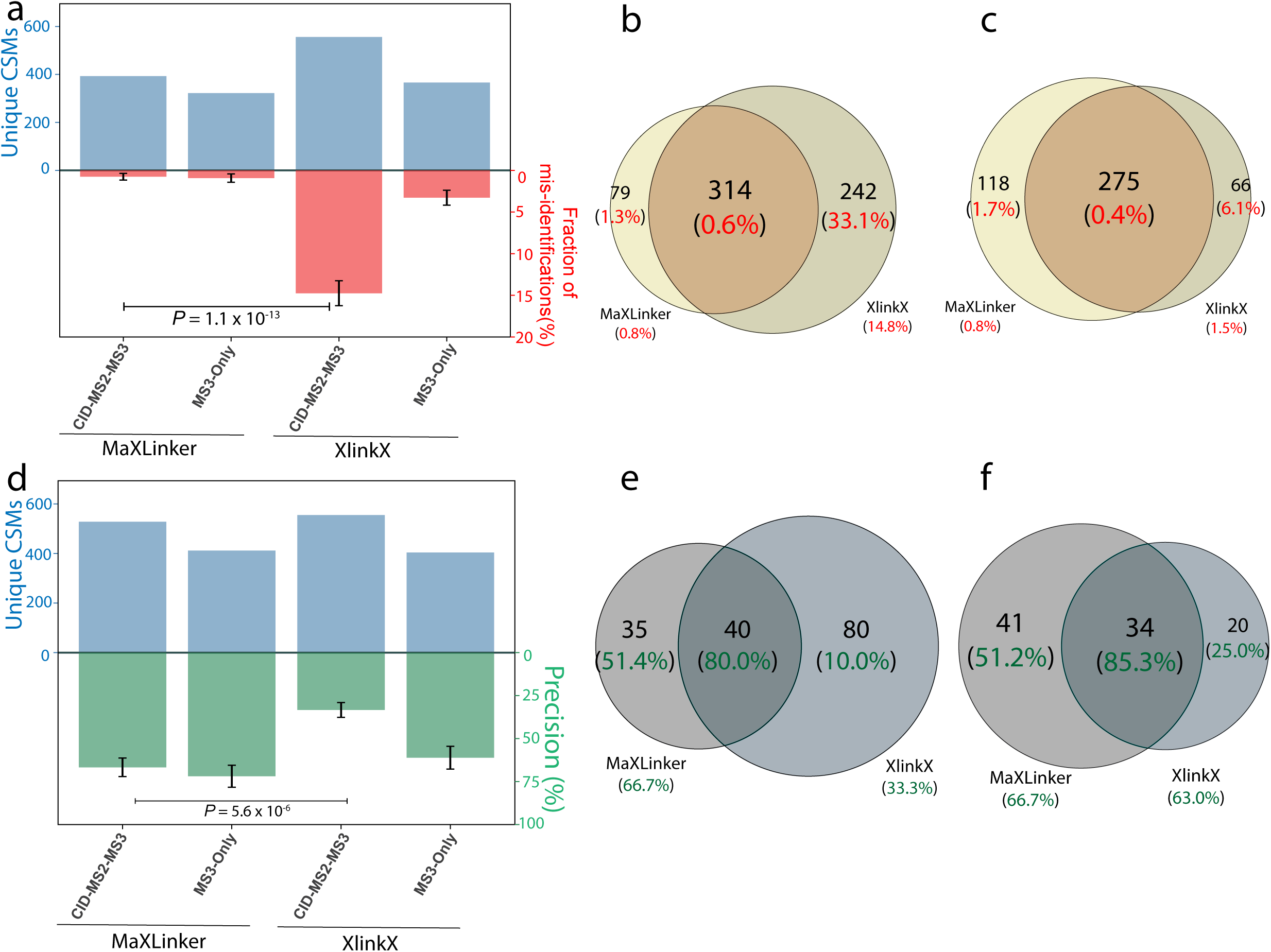
Comparison of MaXLinker’s performance on proteome-wide XL-MS with that of XlinkX. (a) Comparison of the fraction of mis-identifications from MaXLinker and XlinkX at 1% FDR using six *E. coli* MS2-MS3 XL-MS fractions from Liu *et al*^9^. (b) Overlap between CSMs from MaXLinker and XlinkX at 1% FDR showing the respective fraction of mis-identifications in the parentheses. (c) Overlap between CSMs at 1% FDR from MaXLinker and additional filtering on 1% FDR with ‘ΔXlinkX score’ ≥ 50 for XlinkX, showing the respective fraction of mis-identifications in the parentheses. (d) Comparison between MaXLinker and XlinkX in terms of precision using six *E. coli* MS2-MS3 XL-MS fractions from Liu *et al*^9^. (e) Overlap between interprotein CSMs from MaXLinker and XlinkX at 1% FDR showing the respective precision values in the parentheses. (f) Overlap between interprotein CSMs at 1%FDR from MaXLinker and additional filtering on 1% FDR with ‘ΔXlinkX score’ ≥ 50 for XlinkX, showing the respective precision values in the parentheses. (Significance was determined by a two-sided *Z*-test; The error bars represent the estimated standard error of mean).

Our observations provide captivating evidence that, among the three widely used approaches, CID-MS2-MS3 results in cross-links with significantly better quality, most of which rely on MS3 spectra for sequence information. However, the high number of incorrect identifications for CID-MS2-MS3 approach at 1% FDR by XlinkX (the current state-of-the-art search engine) strongly demonstrates the need for an improved search algorithm that can efficiently eliminate false positives while maintaining a minimum number of false negatives.

## MaXLinker: a novel “MS3-centric” approach for cross-link identification

To address the limitations faced by the conventional “MS2-centric” algorithms such as XlinkX for reliable cross-link identifications from MS2-MS3 fragmentation, we designed a novel “MS3-centric” approach (**Fig. 2**). The XlinkX starts the search at MS2-level and attempts to identify CSMs exclusively from the MS2 spectrum, for cases with no available sequence information from MS3-level. However, our analyses revealed that such “MS2-centric” approach could lead up to 14.8% false identifications (**Fig. 1a**). On the contrary, our approach starts the search from MS3-level, which is confirmed through our analyses to be most informative level for the sequences of cross-linked peptides (**Fig. 1**). Additionally, our approach fully utilizes MS2-level to rescue candidate CSMs (‘MS2 Rescue node’) if one of the two cross-linked peptides could be reliably identified from the MS3 spectra (**Fig.2** Node C). Finally, we require all cross-links to match the precursor mass in MS1 (**Fig. 2** Node D) and perform correction for mis-assigned monoisotopic MS1 precursor masses (**Fig.2** Node B). This novel design, where we start with MS3-level information but fully integrate information from both MS2 and MS1 levels, fundamentally enables MaXLinker’s rigorous cross-link identification and validation work-flow.

The general experimental methodology for MS2-MS3 strategy involves precursor selection at multiple stages of mass spectrometry. First, ions above certain threshold charge state (typically ≥ +3 or +4) will be selected for fragmentation at MS2 stage to yield signature ions with predefined mass difference (Δm = 31.97 for DSSO). Further, an iterative search known as ‘targeted inclusion’ is performed by mass spectrometer *on-the-fly* to select ion pairs with signature Δm, following certain prioritization criteria to perform fragmentation at MS3-level to yield two MS3 spectra per peptide in an ideal scenario. MaXLinker accepts ‘.mgf’ files consisting different levels of MS spectra exported using Proteome discover (PD), along with PSM annotations from PD as input (**METHODS**). MaXLinker initiates the search from the MS3-level by performing the mandatory precursor-based mass validation (**Fig. 2** Node ‘A’). Initiating the search from MS3, the most informative level in terms of the peptide sequence information, provides a key advantage to MaXLinker in eliminating potential false positives. If a set of MS3 spectra representing a potential cross-link pass the precursor-based mass validation step (**Fig. 2** Node ‘A’) (Case 1 in **Fig. 2**), it is verified through multiple validation filters (**Fig. 2** Node ‘D’). It is important to note that typically larger size of crosslinked peptides can often result in the mis-assignment of the monoisotopic MS1 precursor mass^15^, thus for cases that fail to pass through the precursor mass-based filter (**Fig. 2** Node ‘A’), MaXLinker inspects the corresponding MS1 spectrum to verify mis-assignment of the monoisotopic MS1 precursor mass (**Fig. 2** Node “B”). Such cases are systematically examined and passed on to the next filter if they satisfy the mass validation step with the adjusted precursor mass. The remaining failed candidates are sent to the ‘MS2 Rescue Module (**Fig. 2** Node ‘C’).

MS2 Rescue Module is another important and unique feature of MaXLinker. As mentioned earlier, this module is triggered if the candidate spectra failed to pass the precursor-based mass validation step (Fig. 2 Node ‘A’) and could not be validated through precursor mass re-assignment. We found that failure to pass these filters often coincided with poor or “uninformative” MS3 spectral data for one of the cross-linked peptides (case 2 in **Fig. 2**). In this case, considering a scenario where the mass spectrometer picked an incorrect Δm pair from the MS2-level having the signature just by chance, MaXLinker attempts to obtain sequence information for the peptide by utilizing fragment ions from the corresponding MS2 spectrum (**Fig. 2** Node C). First, precursor masses for the peptide with poor MS3 spectra are derived using MS2 precursor mass and MS3 precursor masses of the “informative” MS3 spectra (with account for the linker long and short arm modifications) (**Supplementary Figure 5**). An additional validation search is performed on the ions of the corresponding MS2 spectrum to confirm presence of the derived MS3 precursor masses. Subsequently, a PSM search is performed on the deconvoluted MS2 spectrum with the derived masses (both long and short) as the precursor mass. If the search returns at least one reliable PSM, the cross-link candidate (along with sequence information for the ‘rescued’ peptide) is directed to the general validation pipeline for further evaluation (**Fig. 2** Node D). Additionally, the MS2 Rescue module also accounts for cases where the mass spectrometer selects two pairs with signature Δm for MS3, however both pairs represent different charge states of one of the two cross-linked peptides (**Supplementary Figure 6**). Upon completion of the search, a unique list of cross-links is obtained by merging the redundant CSM entries, and a confidence score is assigned to each identification (equation 2 in **METHODS**). Finally, a target-decoy strategy is employed to establish the FDR.

## MaXLinker significantly outperforms XlinkX in both specificity and sensitivity

We evaluated the performance of MaXLinker utilizing MS2-MS3 XL-MS raw files for six *E. coli* fractions from Liu *et al*^9^. First, we utilized the strategy employed in **Fig. 1a** and performed the search using MaXLinker at 1% FDR. We noted that the fraction of mis-identifications was less than 1% (**Supplementary Table 1**), and for majority of the identifications (~82%), the peptide sequence information was derived from MS3 spectra (**Supplementary Table 2**), which agrees with MaXLinker’s fundamental algorithmic design. Next, we compared the results with CSMs identified using XlinkX at 1% FDR on the same set of raw files (**Fig. 3a**). Our analysis showed that MaXLinker evidently outperforms XlinkX, indicated by the extremely significant difference (18-fold lower) in the fraction of mis-identifications (i.e. non-*E. coli* CSMs). We then examined the overlap between identifications from the two search engines (**Fig. 3b**). It clearly reveals that the overlapping fraction from XlinkX has only 0.6% mis-identifications, whereas the non-overlapping CSMs which were identified exclusively by XlinkX contained a large fraction (33.1%) of mis-identifications. Further, using precision as a complementary quality metric, we observed similar results (**Fig. 3d, 3e**). When we repeated the quality analyses by filtering the identifications from XlinkX at different ‘Δ XlinkX score’ cutoffs, we observed that MaXLinker consistently finds 13-31% more cross-links than XlinkX at comparable quality (**Supplementary Figure 4**). Importantly, the CSMs identified exclusively by MaXLinker are of three-fold higher quality than the exclusive identifications by XlinkX, even at the highly stringent cutoff ‘Δ XlinkX score’ ≥ 50 (**Fig. 3c, 3f**). All these results demonstrate that MaXLinker outperforms XlinkX for CSM identifications in both specificity and sensitivity.

**Figure 4.**
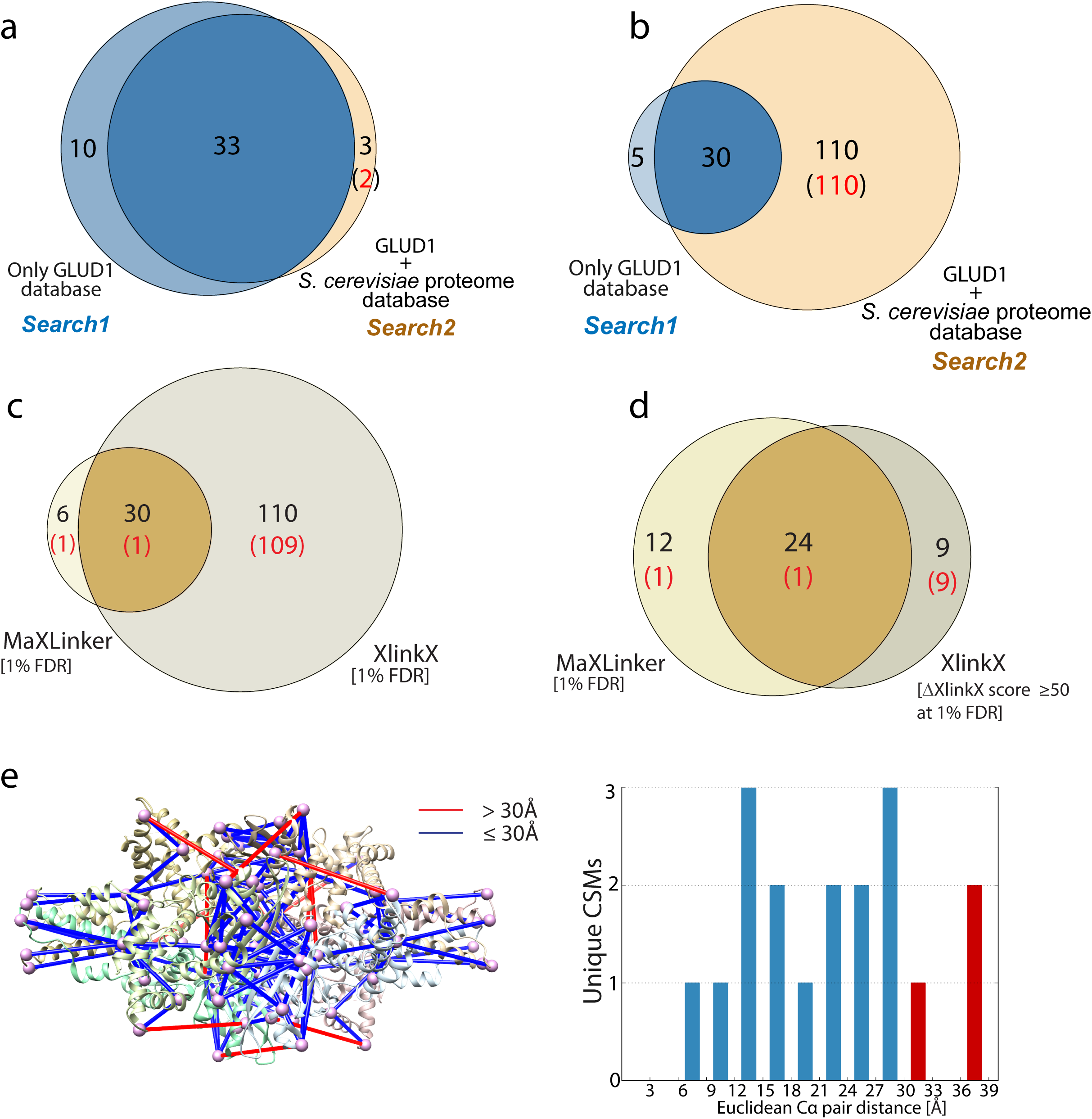
Validation and Comparison of MaXLinker’s performance with that of XlinkX using bovine GLUD1 XL-MS. *Search1* was performed using sequence of only GLUD1 protein as the search database and *Search2* was performed using a concatenated database consisting sequence for GLUD1 along with the entire *S. cerevisiae* proteome. (a) Overlap between MaXLinker’s identifications from *Search1* and *Search2* at 1% FDR. (b) Overlap between identifications from XlinkX from *Search1* and *Search2* at 1% FDR. (c) Overlap of *Search2* identifications at 1% FDR from MaXLinker and XlinkX. (d) Overlap of *Search2* identifications from MaXLinker at 1% FDR and with additional filtering (‘ΔXlinkX Score’ ≥ 50) at 1% FDR from XlinkX. (e) Validation of cross-links from GLUD1 identified using MaXLinker, by mapping them onto its three-dimensional structure (PDB: 5K12). Cross-links exceeding theoretical distance constraint for DSSO (30Å) is shown in red. The histogram shows distance distribution for all the cross-links mapped on to the structure (cross-links with distance >30Å shown in red). The structure mappings were performed using Xlink Analyzer^17^ implemented in UCSF Chimera^18^.

Next, we cross-linked commercially available Bovine Glutamate Dehydrogenase 1 (GLUD1) using DSSO and performed a CID-MS2-HCD-MS3 experiment in our own lab (**METHODS**). We employed MaXLinker to perform two individual CSM searches, *search1*: using Bovine GLUD1 sequence as the search database yielding 43 unique CSMs, and *search2*: with a concatenated database with Bovine GLUD1 and a full proteome of *Saccharomyces cerevisiae*, yielding 36 unique CSMs. We then examined the overlap between CSMs from *search1* and *search2* to inspect MaXLinker’s ability to find true CSMs from single protein in a plethora of false search space. Strikingly, we observed that 33 of 36 (92%) CSMs from *search2* were overlapping with the ones from *search1* (**Fig. 4a**). Out of the remaining 3 CSMs, 2 were mis-identifications, having one of the peptides in the pair from *S. cerevisiae* proteome (false search space). Of note, 10 CSMs were identified exclusively in *search1*. Upon close examination, we noted that MaXLinker rejected those 10 CSM candidates due to either (i) its stringent validation filters or (ii) lower confidence in their PSM assignments, attributable to the drastic increase in the number of competing candidate peptides for individual spectra. On the other hand, when we performed similar analysis using XlinkX, *search1* and *search2* yielded 35 and 140 unique CSMs, respectively. Out of the 140 CSMs from *search2*, 30 were overlapping with *search1* and the remaining 110 had at least one of the peptides from *S. cerevisiae* proteome (mis-identifications) (**Fig. 4b**). We examined the overlap between *search2* identifications from MaXLinker and XlinkX and observed that most of the mis-identifications from XlinkX (109 of 110) were not found by MaXLinker (**Fig. 4c**). Further, we filtered CSMs from XlinkX using ‘Δ XlinkX score’ ≥ 50 and re-inspected the overlap with MaXLinker’s identifications. This filtering step resulted in drastic elimination of false positives (**Fig. 4d**). However, all the non-overlapping CSMs from XlinkX were observed to be mis-identifications. On the other hand, MaXLinker identified 12 CSMs (containing 11 true CSMs) that were missed by XlinkX. For further validation of MaXLinker’s identifications, we mapped CSMs from *search1* on to a three-dimensional structure (**Fig. 4e**) of Bovine GLUD1. We observed that 15 of the 18 mapped CSMs were within the theoretical distance constraint (30Å), and the remaining three CSMs were within 38Å, validating reliable quality of our identifications. This analysis serves as a revealing case study for MaXLinker’s unique ability to identify cross-links with high sensitivity and specificity.

## Our proteome-wide K562 XL-MS study unveils the largest single set of cross-links

Having established the MaXLinker software and optimized the experimental pipeline in our lab, we carried out a comprehensive proteome-wide XL-MS study on human K562 cell lysates, using the CID-MS2-HCD-MS3 strategy. Previous proteome-wide XL-MS studies implemented the strong cation exchange chromatography (SCX) for pre-fractionation of crosslinked proteome samples. Here, to capture a more comprehensive set of cross-links, we employed both SCX and hydrophilic interaction chromatography (HILIC) for our proteome-wide XL-MS study. We then employed MaXLinker for cross-link identification. Our study yielded 9,319 unique cross-links (8,051 intraprotein and 1,268 interprotein with 74.2% precision) at 1% FDR (**Supplementary Table 3**), ~ 3-fold more number of cross-links than that of the latest human proteome XL-MS study^9^. To validate the identified cross-links utilizing available three-dimensional structures, we mapped cross-links from 26S proteasome, which is a large biological complex, on to its three-dimensional structure (**Fig. 5a, 5b**). Out of the 100 cross-links mapped to the structure, 90 were within the theoretical constraint i.e., 30Å. Additionally, we could validate one cross-link that was exceeding 30Å, utilizing a different structure (**Fig. 5c**), suggesting potential conformational changes in the corresponding subunits. Six out of the remaining nine cross-links were within 35Å, and all the others were within 50Å, demonstrating high quality of our identifications. Additionally, interprotein cross-links identified at 1% FDR in our study represent 160 unambiguous novel interactions (**Fig. 5d** and **Supplementary Table 4**).

**Figure 5.**
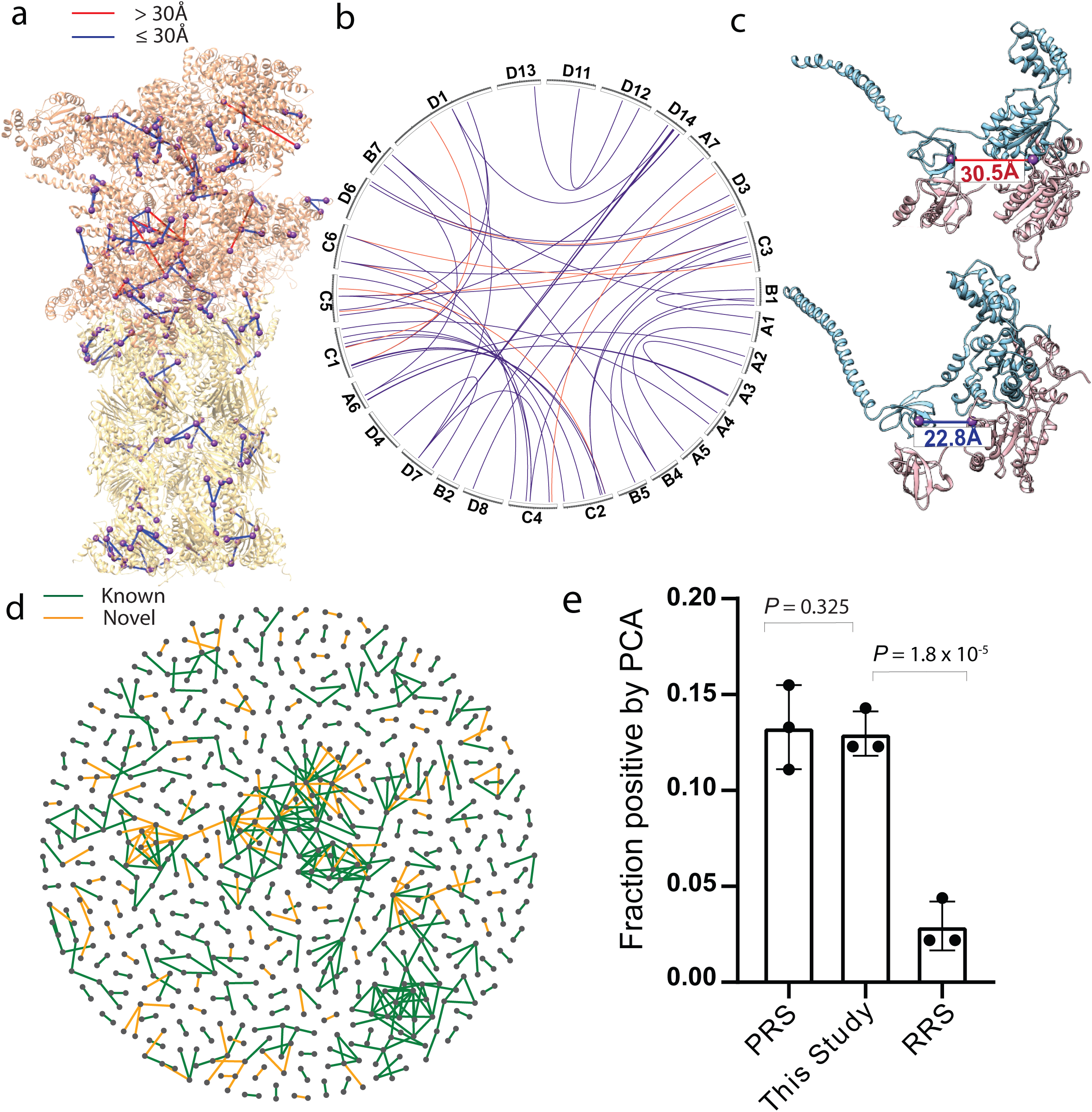
Validation of cross-links and novel interactions identified in the proteome-wide human K562 XL-MS study at 1% FDR. (a) Mapping cross-links from 26S proteasome complex on to a recently published structure (PDB: 5GJQ; cross-links exceeding maximum theoretical constraint 30Å are shown in red). (b) A circular plot showing interprotein cross-links between various subunits from 26S proteasomal complex.(cross-links exceeding maximum theoretical constraint 30Å are shown in red; the plot was generated using Circos^19^.) (c) Validation of a cross-link from 26S proteasome that violate distance constraints (>30Å) in one structure (PDB: 5GJQ) and obey in a different structure (PDB: 5T0J), suggesting potential conformational changes. (d) Network map showing protein-protein interactions identified in the current study. (known interactions are shown in green and novel interactions are shown in orange). (e) Experimental validation of a representative set of 49 novel interactions identified in the current study using Protein-fragment complementation assay (PCA) (mean fraction positive: 0.130) (PRS: Positive Reference Set (45 interactions; mean fraction positive: 0.133); RRS: Random Reference Set (45 interactions; mean fraction positive: 0.029); The error bars represent the standard deviation; Significance was determined by a one-sided Welch Two Sample t-test; 95% confidence interval; *t-statistic* 0.53 for “PRS -This Study”, and 164.75 for “RRS -This Study”; 2 degrees of freedom).

### Systematic experimental validation of novel interactions from our proteome-wide XL-MS study

Furthermore, in order to validate those novel interactions using an orthogonal experimental methodology, a representative subset of them (49 randomly-chosen interactions) was tested individually using a Protein Complementation Assay (PCA). The fraction of PCA-positive interactions among the novel interactions identified in our XL-MS study is statistically indistinguishable (*P* = 0.325) from that of the positive reference set containing well-established interactions in the literature, but significantly higher (*P* = 1.8 × 10^−5^) than that of a negative reference set containing random protein pairs **(Fig. 5e)**^16^. This large set of experimental results demonstrate the high quality of the novel cross-links and corresponding interactions identified in our proteome-wide XL-MS study, and further confirm the reliability and accuracy of MaXLinker.

## DISCUSSION

Machine learning approaches have been an integral part of conventional mass spectrometry-based methods^13^. Here, we extended their applications for comparative quality assessment among multiple proteome-wide XL-MS data sets. In addition to using a false search space from an un-related organism, we demonstrated precision as an effective additional metric for comparative quality assessments. It should be noted that, because a large fraction of true protein interactions is yet to be discovered, precision should not be used as an absolute measure for data quality. Nevertheless, it is an orthogonal and reliable quality metric for comparative assessments of proteome-wide XL-MS studies.

Our systematic analyses revealed for the first time, the limitations of current quality assessment strategies and the drawbacks of the conventional “MS2-centric” cross-link identification approach resulting in high false positive rates (~15%). Our analyses also revealed that for MS2-MS3 strategy, the MS3-level provides sequence information with significantly higher quality when compared to that of the MS2-level, and identification of cross-links exclusively from MS2-level could result in alarmingly high false positive rate. To address these issues, we designed and implemented a novel “MS3-centric” approach (MaXLinker) (**Fig. 2**). The conventional “MS2-centric” methods such as XlinkX start the search from the MS2-level and attempts cross-link identifications without any information from MS3-level, resulting in high fraction of false positives. On the contrary, MaXLinker starts the search from MS3-level and discards any cross-link candidate without reliable sequence information from MS3-level for at least one of the two cross-linked peptides. Furthermore, the “MS2-Rescue” module, along with other novel features such as the correction for mis-assigned MS1 monoisotopic mass (**Fig. 2**), play a crucial role in MaXLinker’s superior sensitivity over the conventional approach, without compromising on the specificity. Overall, MaXLinker significantly outperformed XlinkX with 18-fold lower false positive rate and up to 31% higher number of identifications.

Having MaXLinker in hand, we reported the largest single data set from proteome-wide XL-MS consisting 9,319 cross-links at 1% FDR, representing 160 unambiguous novel interactions. Moreover, to our knowledge, this is the first study that performed a large-scale orthogonal experimental validation of novel interactions identified from a proteome-wide XL-MS study.

With the constant technical advancements in XL-MS methodologies, reliable search algorithms such as MaXLinker will play a highly significant role in the success of future cross-linking studies. Moreover, the expanding size of cross-link datasets would allow researchers to investigate interaction networks in many disease phenotypes more thoroughly, thereby enabling us to better understand the underlying molecular mechanisms.

## ACKNOWLEDGEMENTS

We thank Rosa Viner for support in data processing with XlinkX work-flow in Proteome Discoverer and Shayne Wierbowski for helpful suggestions regarding data representation. We thank Robert Fragoza for the assistance with PCA experiments. We thank Elizabeth Anderson and Robert Sherwood for their technical support in sample preparation.

## AUTHOR CONTRIBUTIONS

H.Y. conceived and oversaw all aspects of the study. K.Y. and H. Y. designed and implemented MaXLinker algorithm with assistance from IM and MCL. A.K.L. developed the user interface for MaXLinker. K.Y. performed the computational analyses. T.W., J.L., and E.S. performed laboratory experiments. K.Y. wrote the manuscript with inputs from T.W., M.C.L., M.B.S., S.Z., and H.Y.

### COMPETING FINANCIAL INTERESTS

The authors declare no competing financial interests.

## METHODS

### Cell culture and whole cell lysate preparation

The K562 cells (ATCC^®^ CCL-243™) were purchased from American Type Culture Collection (ATCC). The cells were maintained in the Iscove’s Modified Dulbecco’s Medium (IMDM) (ATCC) supplemented with 10% fetal bovine serum (FBS) (ATCC) at 37°C with humidified ambient a™osphere containing 5% CO_2_. The K562 cells were collected and washed three times with cold PBS. The cells were then resuspended in cold buffer composed of 50 mM HEPES, 150 mM NaCl, pH 7.5 supplemented with Protease Inhibitor Cocktail (Roche). The resuspended cells were lysed on ice by sonication (Amplitude 10% for 5 sec and repeat 6 times), followed by centrifugation at 15,000 *g* for 10 min at 4°C. The supernatant was collected and measured the protein concentration using Bio-Rad Protein Assay Dye (Bio-Rad).

### Cross-linking of bovine glutamate dehydrogenase (GDH) and human proteome

DSSO (Thermo Fisher Scientific) was freshly prepared as a 50 mM stock solution by dissolving in anhydrous DMSO. The 1 mg/mL pure bovine glutamate dehydrogenase (GDH) protein (Sigma) was reacted with 1 mM DSSO in 50 mM HEPES buffer, 150 mM NaCl, pH 7.5 for 30 min at room temperature. Similarly, the 1 mg/mL lysate of K562 cells were incubated with 1 mM DSSO for 1 hour at room temperature. Both cross-linking reactions were terminated by 50 mM Tris-Cl buffer, pH 7.5.

### Processing of DSSO-cross-linked samples for analysis

The DSSO-treated protein samples were processed as previously described^20, 21^. Briefly, the cross-linked GDH was denatured in 1% SDS, reduced by DTT, and alkylated with iodoacetamide, followed by precipitated in cold acetone-ethanol solution (acetone:ethanol:acetic acid=50:49.9:0.1, v/v/v). The precipitates were dissolved in 50 mM Tris-Cl, 150 mM NaCl, 2 M urea, pH 8.0 and digested by Trypsin Gold (Promega) at 37°C overnight. After digestion, the sample was acidified by 2% trifluoroacetic acid-formic acid solution, desalted through Sep-Pak C18 cartridge (Waters), and dried using SpeedVac™ Concentrator (Thermo Fisher Scientific). The sample was then reconstituted in 0.1% trifluoroacetic acid and stored in -80°C before mass spectrometry analysis. The DSSO-cross-linked human proteome was processed identically as described above except that the TPCK-treated trypsin was used for digestion and the sample was stored after drying.

### Fractionation by Strong Cation Exchange (SCX)

The SCX fractionation was performed on a Dionex UltiMate 3000 Series instrument (Thermo Fisher Scientific) using a PolySULFOETHYL A column (5 µm, 200 Å, 2.1 x 200 mm; PolyLC) with 10 mM potassium phosphate monobasic in 25% acetonitrile, pH 3.0 as Buffer A and 10 mM potassium phosphate monobasic/500 mM potassium chloride in 25% acetonitrile, pH 3.0 as Buffer B. All eluents were filtered through a 0.22 µm Durapore membrane (EMD Millipore Corporation) and stored at 4°C until use. Prior to injection, the 1 mg of trypsin-digested sample was reconstituted in 25% acetonitrile/0.1% formic acid (v/v) and filtered through a Spin-X centrifuge tube filters (cellulose acetate membrane, 0.22 µm; Corning) by following manufacturer’s recommended protocol. The fractionation was performed at a flow rate of 200 µL/min using a linear gradient from 5-60% of Buffer B in 40 min and 60-100% of Buffer B in an additional 10 min. A total of 60 fractions were collected using a 96-well plate at 1-min intervals monitored by the absorbance at 220 nm and 280 nm. The fractions collected from 23 to 60 min were desalted using SOLA HRP SPE cartridges (Thermo Scientific). The eluted peptides were dried by speed vacuum and stored at -20°C until LC-MS analysis.

### Fractionation of cross-linked peptides by hydrophilic interaction liquid chromatography (HILIC)

The DSSO-cross-linked human peptides in 70% acetonitrile and 1% formic acid were fractionated and enriched by hydrophilic interaction liquid chromatography (HILIC). The HILIC fractionation was performed on a Dionex UltiMate 3000 Series instrument (Thermo Fisher Scientific) equipped with a TSKgel Amide-80 column (3 µm, 4.6 mm x 15 cm; Tosoh). The three following solvents were used: 90% acetonitrile (solvent A), 80% acetonitrile and 0.005% trifluoroacetic acid (solvent B), 0.025% trifluoroacetic acid (solvent C). All the runs were performed at a flow rate of 600 µl/min using the following gradients: 0-5 min (0-98% B and 0-2% C); 5-55 min (98-75% B and 2-25% C); and 55-60 min (75-5% B and 25-95% C). The fractions were collected per 30 seconds. Each fraction was dried and stored at -80°C for further analysis.

### LC-MS^n^ analysis

The HILIC fractions were reconstituted in 0.1% trifluoroacetic acid. The samples were analyzed using an EASY-nLC 1200 system (Thermo Fisher Scientific) equipped with an 125-µm x 25-cm capillary column in-house packed with 3-µm C18 resin (Michrom BioResources) and coupled online to an Orbitrap Fusion Lumos Tribrid mass spectrometer (Thermo Fisher Scientific). The LC analysis were performed using the linear gradients of solvent A composed of 0.1% formic acid and solvent B composed of 80% acetonitrile and 0.1% formic acid with a total run time of 180 min at a flow rate of 300 nl/min. For MS^n^ data acquisition, the CID-MS2-HCD-MS3 method was used. Briefly, the MS^1^ precursors were detected in Orbitrap mass analyzer (375-1500 m/z, resolution of 60,000). The precursor ions with the charge of 4+ to 8+ were selected for MS^2^ analysis in Orbitrap mass analyzer (resolution of 30,000) with the collision energy of collision-induced dissociation (CID) at 25%. The peaks with a mass difference of 31.9721 Da, which is a signature of cleaved DSSO-cross-linked peptides, in CID-MS2 spectra were selected for further MS^3^ analysis. The selected ions were fragmented in IonTrap using higher-energy collisional dissociation (HCD) with the collision energy at 35%.

### Validation of newly identified protein-protein interactions by protein complementation assay (PCA)

The ORFs of a total of 49 protein pairs in pDONR223 plasmid were picked from hORFeome v8.1 library^22^. The bait and prey protein of each protein pair was cloned into the expression plasmids containing the complementation fragments of a fluorescent protein Venus using Gateway LR reactions. The success of the LR reactions with desired ORF was confirmed by PCR using the plasmid-specific primers. To perform PCA, HEK293T cells were cultured in Dulbecco’s Modified Eagle Medium (DMEM) supplemented with 10% fetal bovine serum (FBS) (ATCC) in black 96-well flat-bottom plates (Costar) with 5% CO2 at 37°C. At 60-70% confluency, the cells were co-transfected with the plasmids containing the bait and prey ORF (100 ng for each) pre-mixed with polyethylenimine (PEI) (Polysciences Inc.) and OptiMEM (Gibco). A total of 49 bait and prey ORF pairs along with previously published 45 positive reference pairs and 45 negative reference pairs were examined and distributed across different plates^23, 24^. After 68 hours, the fluorescence of the transfected cells was measured using Infinite M1000 microplate reader (Tecan) (excitation= 514 ± 5 nm / emission = 527 ± 5 nm). The PCA experiments were performed and analyzed in triplicate. The p-values were calculated using a paired one-tailed t-test.

### Precision

Precision for proteome-wide XL-MS studies can be defined as the fraction of the identified interprotein cross-links from previously known protein-protein interactions. It can be derived using the following equation:

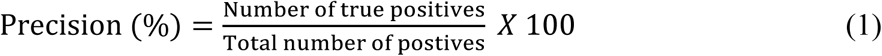

where, ‘positives’ include all the identified interprotein cross-links, and ‘true positives’ refer to cross-links from known protein-protein interactions. We compiled the known protein-protein interactions for *E. coli* and *H. sapiens* from seven primary interaction databases. These databases include IMEx^25^ partners IntAct^26^, MINT^27^, and DIP^28^; IMEx observer BioGRID^29^; and additional sources HPRD^30^, MIPS^31^, and iRefWeb^32^. Furthermore, iRefWeb combines interaction data from CORUM^33^, BIND^34^, MPPI^31^ and OPHID^35^.

### Data processing

The raw data files were converted, and the spectra were exported as ‘.mgf’ (MS1 spectra as ‘.dta’) files using Proteome Discoverer 2.1 software (PD 2.1). SEQUEST searches were performed using PD 2.1 with the following settings: *precursor mass tolerance*: 20ppm (10 ppm for *MS2 rescue module*); *MS3 fragment ion mass tolerance*: 0.6 Da (0.05 Da for *MS2 rescue module*); *fixed modification*: Cys carbamidomethylation; *variable modifications*: Met oxidation, Long arm of DSSO, Short arm of DSSO; *max. equal modification per peptide*: 3; *max. missed cleavages*: 3, *minimum peptide length*: 5. Concatenated target-decoy databases are used for various PSM searches performed during the study. Target sequences were downloaded from uniport database^36^ (with filter ‘reviewed’) and a corresponding decoy database was generated by randomizing the sequences using an in-house python script. ((i) *Escherichia coli*: 5268 sequences; downloaded on 28^th^ October 2017, (ii) *Saccharomyces cerevisiae:* 7904 sequences; downloaded on 28^th^ September 2017, and (iii) *Homo Sapiens*: 42202 sequences; downloaded on 23^rd^ June 2017).

For XlinkX searches, all the raw files were processed using XlinkX v2.0 implemented in Proteome Discoverer software version 2.2 (PD 2.2). PD templates for different XlinkX search methodologies were obtained from Rosa Viner (Thermo fisher Scientific). All the searches were performed at 1% FDR cut-off and the CSMs were exported (after applying filter “*Is Decoy*: False”). For “MS3-Only” category, results from “CID-MS2-MS3” were reprocessed with option “Reprocess: Last Consensus Step” with “Ignore reporter scan: True” in “Xlinkx Crosslink Grouping” node. This set contained a list of all CSMs (includes multiple identifications representing a cross-linked peptide pair). This set of data was used for comparisons shown in Supplementary Fig. 1. Next, Those CSMs for were further processed to obtain a list of unique CSMs (In case of multiple CSMs with different cross-link positions, only one of them was retained to avoid potential biases due to over-representation of certain peptide pairs). The resulting set of CSMs were used for comparisons shown in Fig. 1, Fig. 3a, 3b, 3c, 3d, 3e, 3f, Supplementary Figure 2, Supplementary Figure 3, Supplementary Figure 4. Same procedure was followed to obtain the unique CSMs for GLUD1 analysis shown in Fig. 4b, 4c, and 4d.

### Description of MaXLinker

MaXLinker runs in two steps: (i) *pre-processing* generates a ‘.MS2_rescue.mgf’ file, which is needed for the PSM search in PD 2.1 to be eventually used in the main search. (ii) *cross-link search* accepts .mgf files with different levels of MS spectra(MS1, MS2 and MS3), and two files containing the list of PSMs from PD2.1 SEQUEST search on MS3 spectra and ‘MS2_rescue’ spectra. After the Main MaXLinker search, confidence score is assigned to each cross-link and it is derived using the following equation:

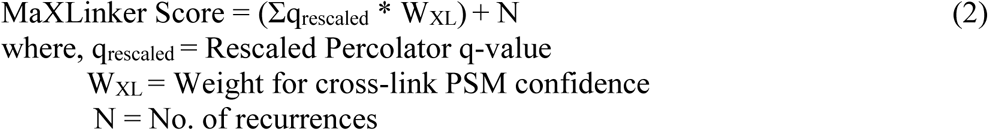

Moreover, MaXLinker utilizes the target-decoy strategy to establish the FDR. A concatenated database consisting target and decoy (random) sequences is used for the PSM search and the FDR is calculated using the equation:

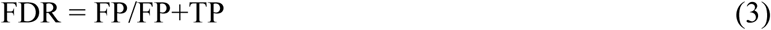

where, FP denote false positive hits and TP denote true positive hits. For cross-link identification, TP represent the number of cross-links with both linked peptides from the target database and FP represent the number of cross-links with at least one of the linked peptides from decoy database.

The identified cross-links were annotated as ‘interprotein’ if neither of the linked peptides were derived from a common protein (with the exception where, both the linked peptides from a common protein, were identical or one of them was a complete subset of the other and the peptide occurs only once in the protein sequence). Cross-links that did not satisfy the aforementioned criteria were annotated as ‘intraprotein’.

### Statistics

Statistical analyses were performed using a two-sided Z test or a one-sided Welch Two Sample t-test, as indicated in the figure captions. Exact *P* values are provided for all compared groups.

### Data availability

All cross-links are reported in the Supplementary information. Additional data that support the findings of this study are available from the corresponding author upon request

