## Supplementary Figures for "MaXLinker: proteome-wide cross-link identifications with high specificity and sensitivity"

**Supplementary Figure 1**. Comparative evaluation of different XL-MS acquisition strategies at the redundant CSM level at 1% FDR. Significance was determined by a two-sided *Z*-test. (analysis at unique CSM level is shown in Fig. 1).

**Supplementary Figure 2**. Quality assessment of cross-links from multiple XL-MS acquisition strategies utilizing false search space (left panel) and precision (right panel). CSMs were identified using XlinkX at 1%FDR with additional filtering cut-offs for 'ΔXlinkX Score'.

**Supplementary Figure 3.** Dissection of CSMs from the ensemble i.e., CID-MS2-MS3-ETD-MS2 approach at 1% FDR in terms of quality of the two inherent approaches (i.e., CID-MS2-MS3 and CID-MS2-ETD-MS2). Venn Diagrams show the overlap of CSMs at 1% FDR, along with further filtering at different 'ΔXlinkX Score' cut-offs. Left panel shows CSM overlap between the two approaches (fraction of mis-identifications is shown in the parentheses) and the right panel shows overlap for the inter-protein cross-links (precision is given in parentheses). XlinkX was used to process the raw files for six *E. coli* CID-MS2-MS3-ETD-MS2 XL-MS fractions from Liu *et al*[9].

**Supplementary Figure 4.** Venn diagrams showing the overlap of CSMs from MaXLinker at 1% FDR with CSMs identified by XlinkX at 1% FDR with further filtering at different 'ΔXlinkX Score' cut-offs for XlinkX (using the same set of raw files for six *E. coli* MS2-MS3 XL-MS fractions from Liu *et al*[9]). Left panel shows results for analysis using false search space (fraction of mis-identifications given in parentheses below the number of CSMs) and the right panel represents precision-based analysis (precision given in parentheses below the number of interprotein CSMs).

**Supplementary Figure 5.** Illustration of MaXLinker's 'MS2 Rescue' module, in a case where the precursor mass validation step fails along with uninformative MS3 spectra for either of the peptides

**Supplementary Figure 6.** Illustration of MaXLinker's 'MS2 Rescue' module, in a case where the precursor mass validation step fails along with all four MS3 spectra potentially representing different charge states of one peptide.

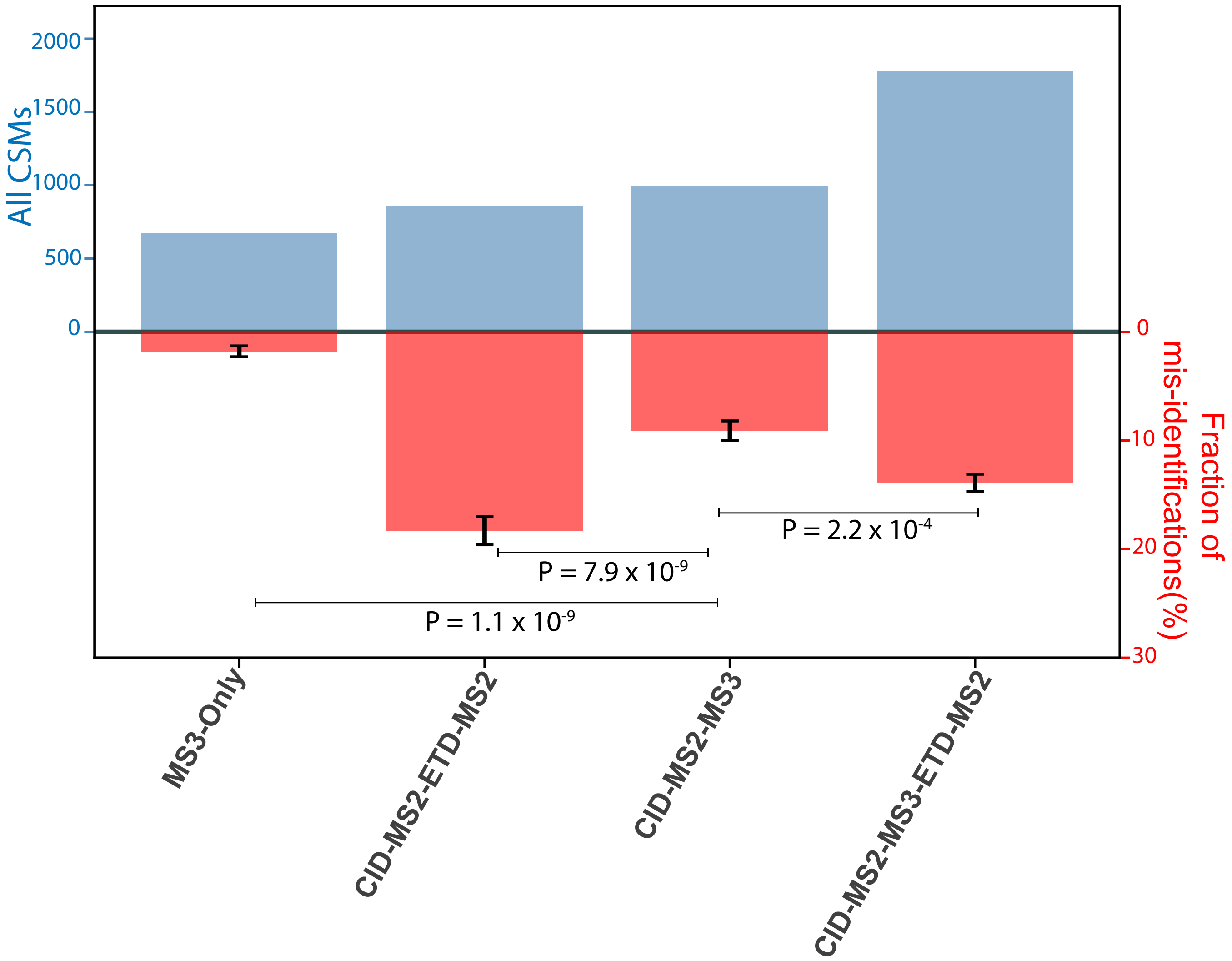


**Supplementary Figure 1**

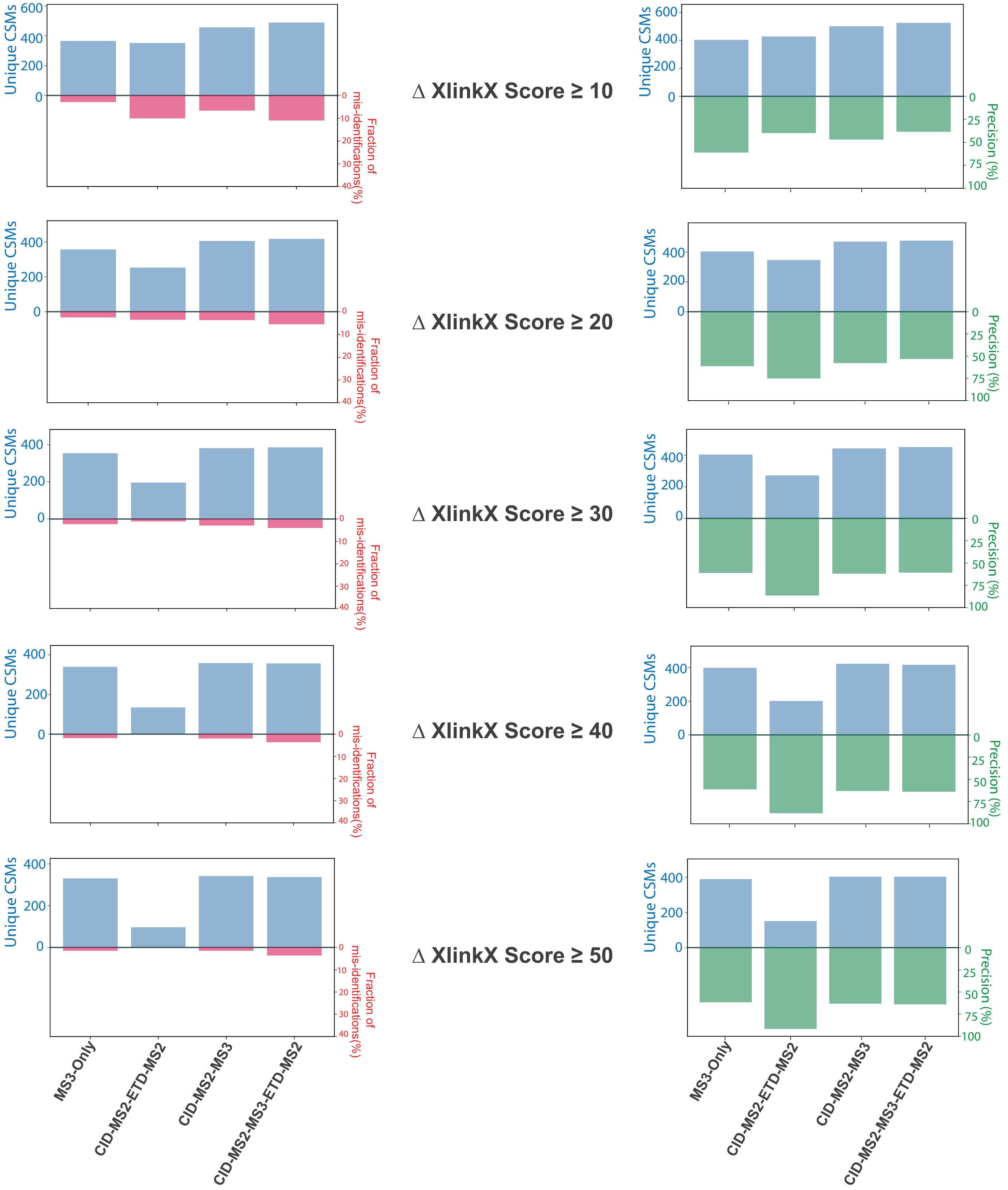


### Supplementary Figure 2

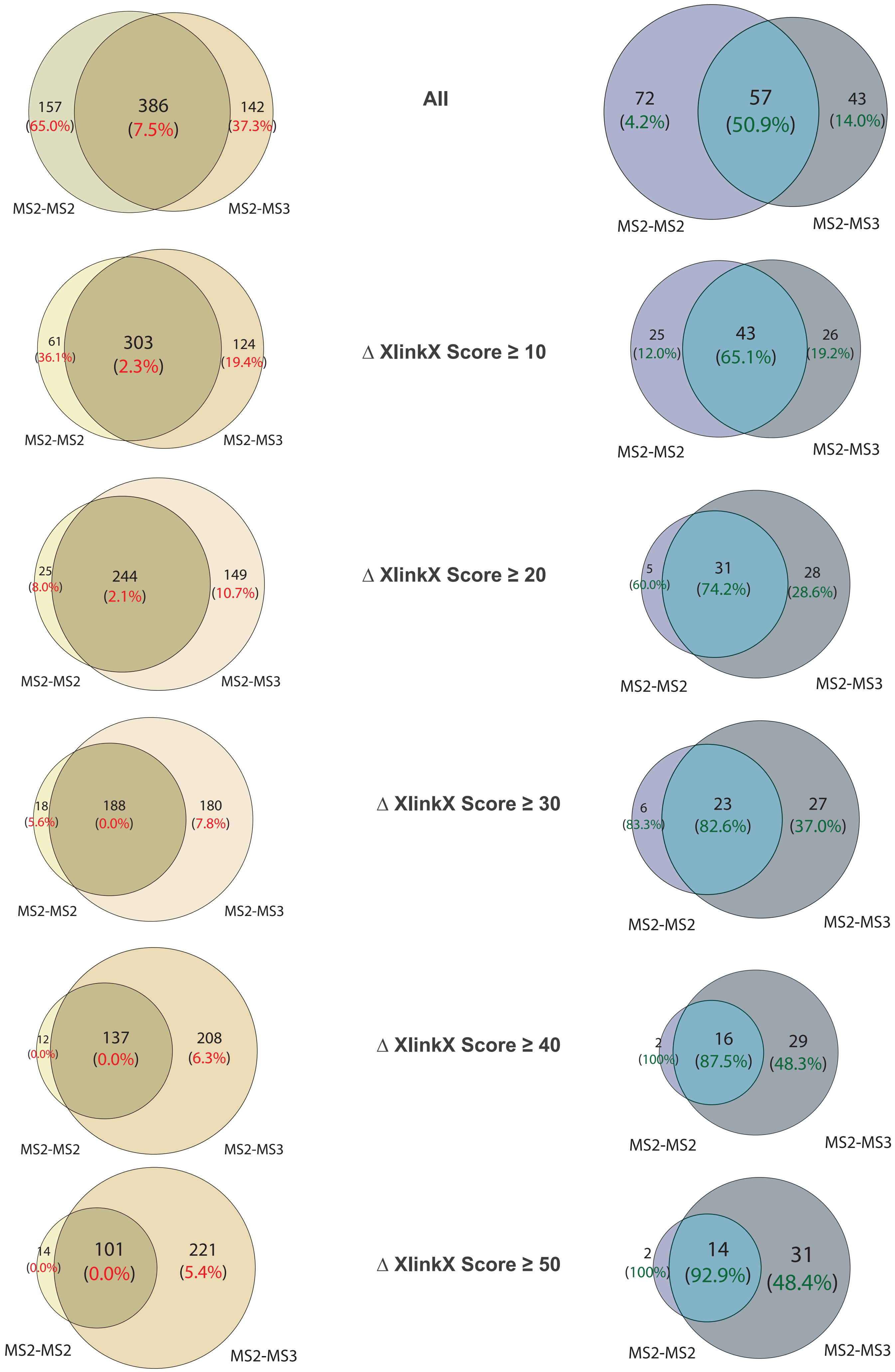


**Supplementary Figure 3**

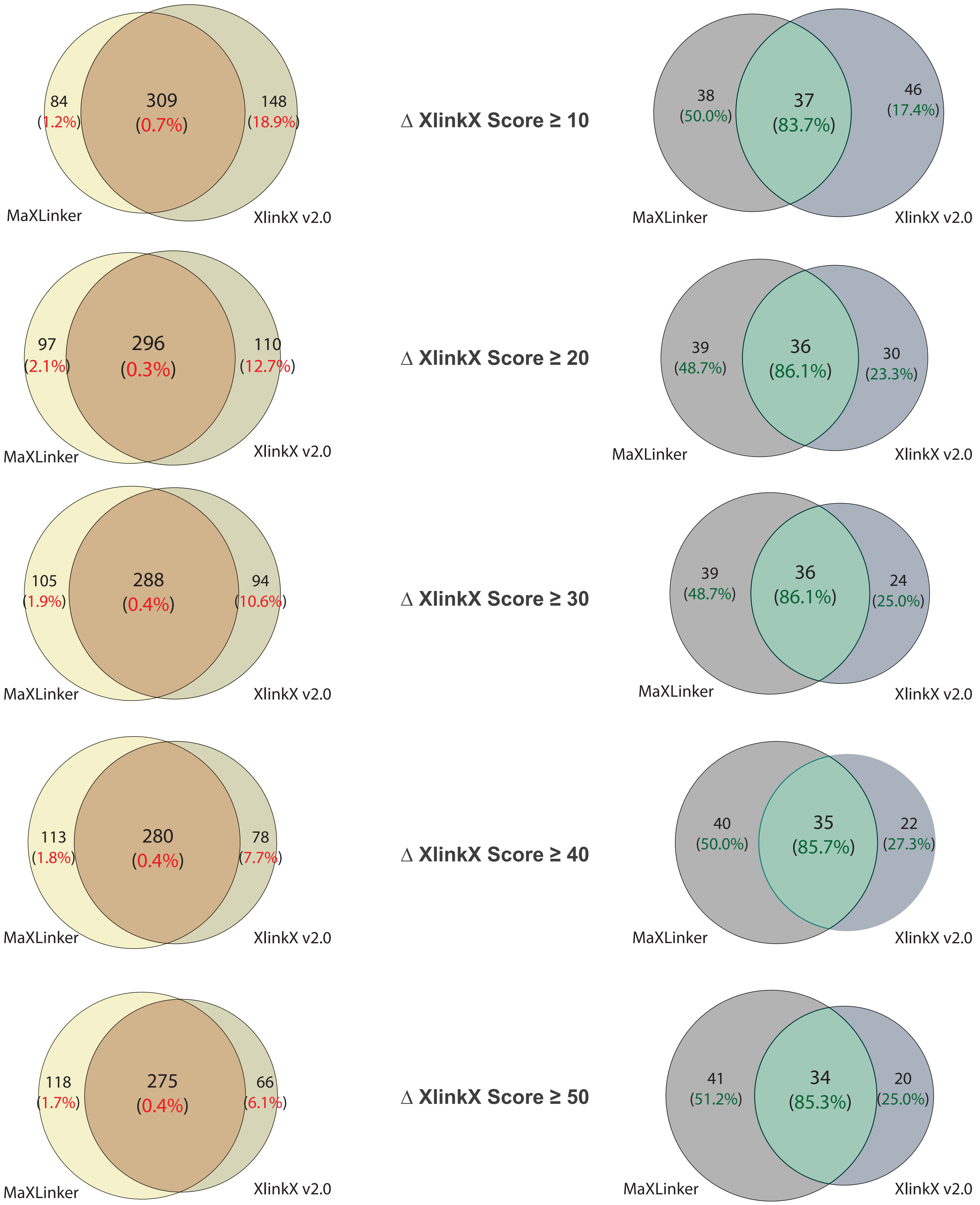


**Supplementary Figure 4**

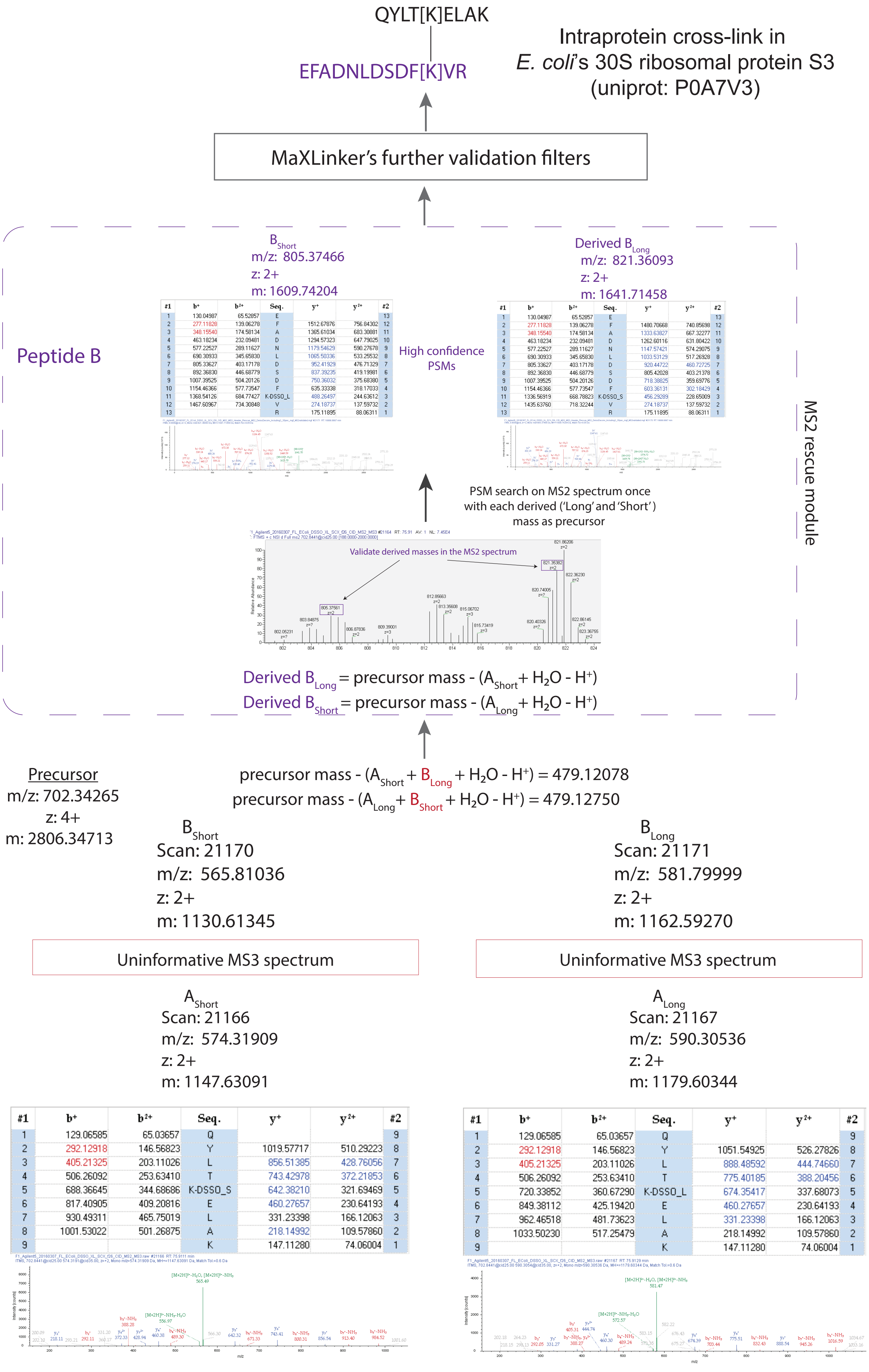


| #1 | b+ | b2+ | Seq. | y+ | y2+ | #2 |
|---|---|---|---|---|---|---|
| 1 | 129.06585 | 65.03657 | Q | | | 9 |
| 2 | 292.12918 | 146.56823 | Y | 1019.57717 | 510.29223 | 8 |
| 3 | 405.21325 | 203.11026 | L | 856.51385 | 428.76056 | 7 |
| 4 | 506.26092 | 253.63410 | T | 743.42978 | 372.21853 | 6 |
| 5 | 688.36645 | 344.68686 | K-DSSO_S | 642.38210 | 321.69469 | 5 |
| 6 | 817.40905 | 409.20816 | E | 460.27657 | 230.64193 | 4 |
| 7 | 930.49311 | 465.75019 | L | 331.23398 | 166.12063 | 3 |
| 8 | 1001.53022 | 501.26875 | A | 218.14992 | 109.57860 | 2 |
| 9 | | | K | 147.11280 | 74.06004 | 1 |

| #1 | b+ | b2+ | Seq. | y+ | y2+ | #2 |
|---|---|---|---|---|---|---|
| 1 | 129.06585 | 65.03657 | Q | | | 9 |
| 2 | 292.12918 | 146.56823 | Y | 1051.54925 | 526.27826 | 8 |
| 3 | 405.21325 | 203.11026 | L | 888.48592 | 444.74660 | 7 |
| 4 | 506.26092 | 253.63410 | T | 775.40185 | 388.20456 | 6 |
| 5 | 720.33852 | 360.67290 | K-DSSO_L | 674.35417 | 337.68073 | 5 |
| 6 | 849.38112 | 425.19420 | E | 460.27657 | 230.64193 | 4 |
| 7 | 962.46518 | 481.73623 | L | 331.23398 | 166.12063 | 3 |
| 8 | 1033.50230 | 517.25479 | A | 218.14992 | 109.57860 | 2 |
| 9 | | | K | 147.11280 | 74.06004 | 1 |

**Supplementary Figure 5**

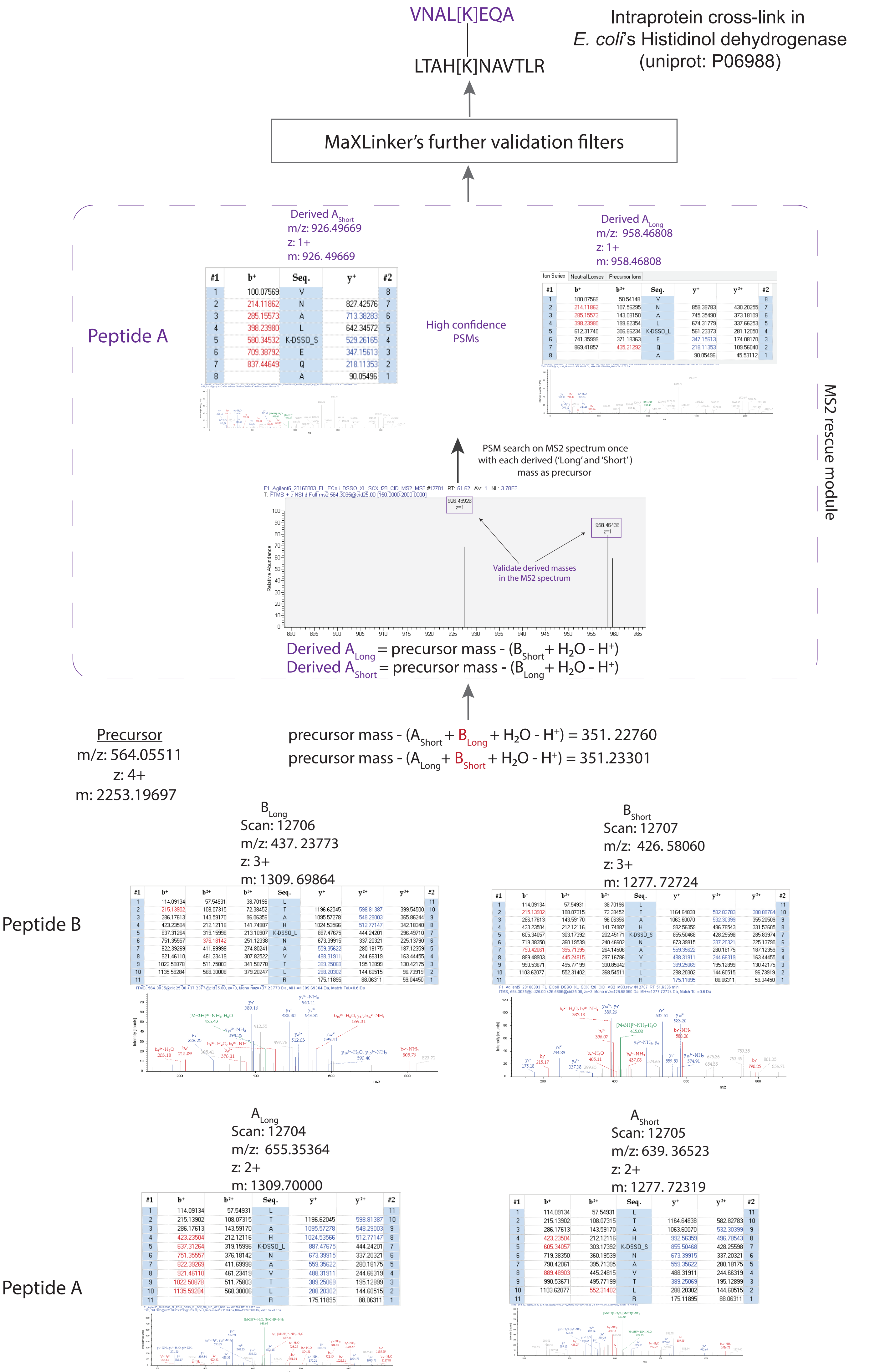


**Supplementary Figure 6**
